# Ion counting demonstrates a high electrostatic potential of the nucleosome

**DOI:** 10.1101/514471

**Authors:** Magdalena Gebala, Stephanie Johnson, Geeta Narlikar, Daniel Herschlag

## Abstract

The fundamental unit of chromatin is the nucleosome, which comprises of DNA wrapped around a histone protein octamer. The association of positively charged histone proteins with negatively charged DNA is intuitively thought to attenuate the electrostatic repulsion of DNA, resulting in a weakly charged nucleosome complex. In contrast, theoretical and computational studies suggest that the nucleosome retains a strong, negative electrostatic field. Despite their fundamental implications for chromatin organization and function, these opposing models have not been experimentally tested. Herein, we directly measure nucleosome electrostatics and find that while nucleosome formation reduces the complex charge by half, the nucleosome nevertheless maintains a strong negative electrostatic field. Further, our results show that the wrapping of DNA around a histone octamer increases the propensity of the DNA to make interactions with multivalent cations like Mg^2+^. These findings indicate that presentation of DNA on a nucleosome may more strongly attract positively-charged DNA binding proteins. Our studies highlight the importance of considering the polyelectrolyte nature of the nucleosome and its impact on processes ranging from factor binding to DNA compaction.

## Introduction

The eukaryotic nuclear DNA forms a highly compact and organized structure referred to as chromatin. Despite this compaction, chromatin is accessible to a vast cohort of macromolecules which regulate its structure, dynamics, and structural plasticity and thereby influence gene expression and determine cell differentiation and state.^1-4^

The most basic level of nuclear DNA compaction is driven by association with positively charged histone proteins to form nucleosomes (Figure 1A). The nucleosome complex is composed of 147 base-paired (bp) DNA wrapped in a left-handed helix with ∼1.7 superhelical turns around the core of eight histone proteins, two copies each of H2A, H2B, H3 and H4.^5-7^ The DNA associates with the histone octamer via backbone and minor groove interactions that involve salt bridges, water-mediated and direct hydrogen bonds, and deep insertions of positively charged arginine into each DNA minor groove facing the central histone octamer.^7-9^

**Figure 1.**
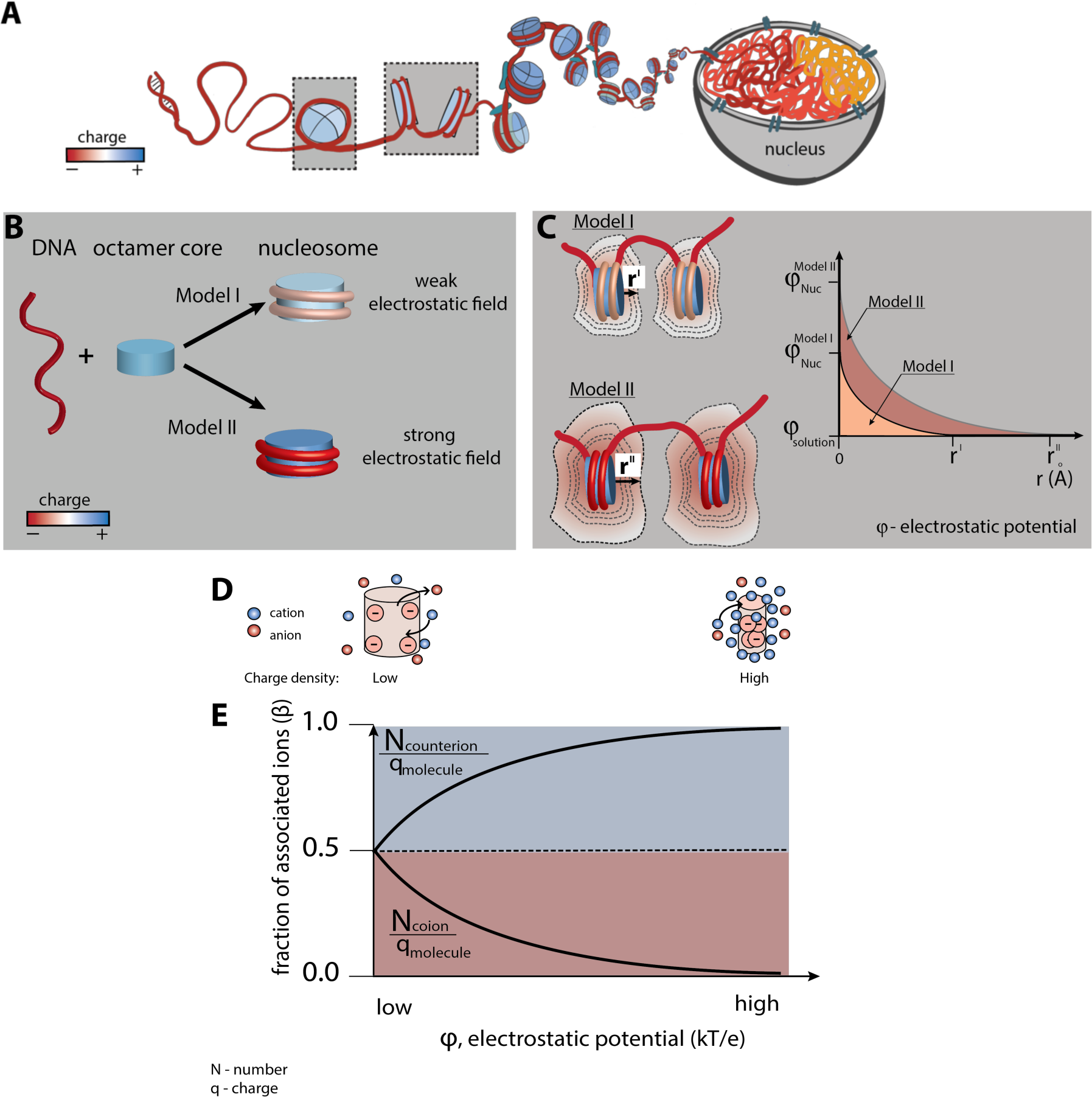
Nucleosome electrostatics: models and implications. (A) Schematic representation of nucleosome formation and DNA compaction. (B) Models of nucleosome electrostatics. According to Model I, the electrostatic field around the nucleosome is weak due to compensatory electrostatic interactions between the DNA and the positively charged histone octamer. In Model II the electrostatic field remains strong, as has been proposed by theoretical and computational studies.^13, 1^^4^ (C) Schematic representation of the effect of nucleosome electrostatics on their propensity to compact according to Models I and II. The distance for full screening of electrostatic repulsion is considerably less for Model I (top) than for Model II (bottom; r^I^ vs. r^II^). (D) Schematic representation of an ion atmosphere around a low charge density (ρ) (left) and high charge density molecule (right). (E) Fraction of associated counterions (e.g., cations around negatively charged molecule) and coions (e.g., anions around negatively charged molecule) within the ion atmosphere around a molecule as a function of its electrostatic potential. The magnitude of the electrostatic potential correlates with the charge density of the molecule: the higher the charge density, the large magnitude of the electrostatic potential and stronger counterion attraction, as depicted by the cartoons in part (D).

DNA is one of the most charged polymers in nature, carrying two negative charges per base pair and generating a strong negative electrostatic field that influences its mechanical properties and its interactions with proteins and small molecules.^10^ This field provides a substantial barrier to DNA compaction in the form of DNA/DNA self-repulsion. It is generally assumed that that association of the DNA around the positively charged histone octamer “can effectively neutralize the negatively charged DNA backbone”, ameliorating repulsive interactions and facilitating compaction to form higher-order nucleosomal structures.^11^ We refer to this as Model I and depict it schematically in Figure 1B.

A contrasting model, Model II in Figure 1B, arises from the fact that the negative charge of 147 bp DNA exceeds the positive charge of the histone octamer by approximately two-fold, resulting in a nucleosome complex that remains highly negatively charged. Theoretical calculations (e.g., Poisson Boltzmann mean-field calculations and all-atom models) that consider the DNA and histone core charges predict that the close wrapping of the DNA in the nucleosome results in enhanced local negative charge density (i.e., ρ, charge per volume) and an overall *increase* in electrostatic field despite the nucleosome’s lower net charge relative to free DNA.^12-14^

These models predict diametrical physicochemical properties and behaviors of nucleosomes, so distinguishing between them is of fundamental importance to understanding DNA compaction and the interactions that regulate chromatin function. While Model I is widely espoused because of its intuitiveness, it has severe limitations. Proteins that interact with DNA to control transcription, repair damage, and remodel chromatin structure often rely on electrostatic attraction and electrostatically-guided one-dimensional diffusion to locate binding sites and to bind DNA.^15-17^ These abilities would be lost if DNA’s electrostatic field were nullified. An attractive feature of Model I is the prediction that charge neutralization lowers repulsion and allows nucleosomes to approach more closely. However, nuclear DNA is considerably more compact than predicted from a simple absence of repulsion.^12, 18-21^, and Model I predicts weakening of DNA-protein interactions in nucleosomes that might otherwise be responsible for further compaction. In contrast, Model II predicts that the nucleosomal DNA maintains a strong electrostatic attraction for proteins and cellular polycations, allowing these and other favorable interactions that may be required to bridge nucleosomes and further compact DNA.

Given these stark differences, it is important to experimentally test these models. “Ion counting” is arguably the most effective experimental approach to analyze nucleic acid electrostatics and test theoretical predictions.^22-27^ It uses equilibration with a buffer solution followed by inductively coupled plasma mass spectroscopy (BE-MS) to precisely determine the number of ions that interact with a nucleic acid and form an ion atmosphere around the molecule—i.e., the number of cations that are attracted to and anions that repelled from the DNA over those present in bulk (Figure 2). These numbers are directly related to the magnitude of a molecule’s electrostatic field and can thus be used to infer the strength of electrostatics interactions (see “Strategy to measure the electrostatics of nucleosomes”).^22-25, 28-36^

**Figure 2.**
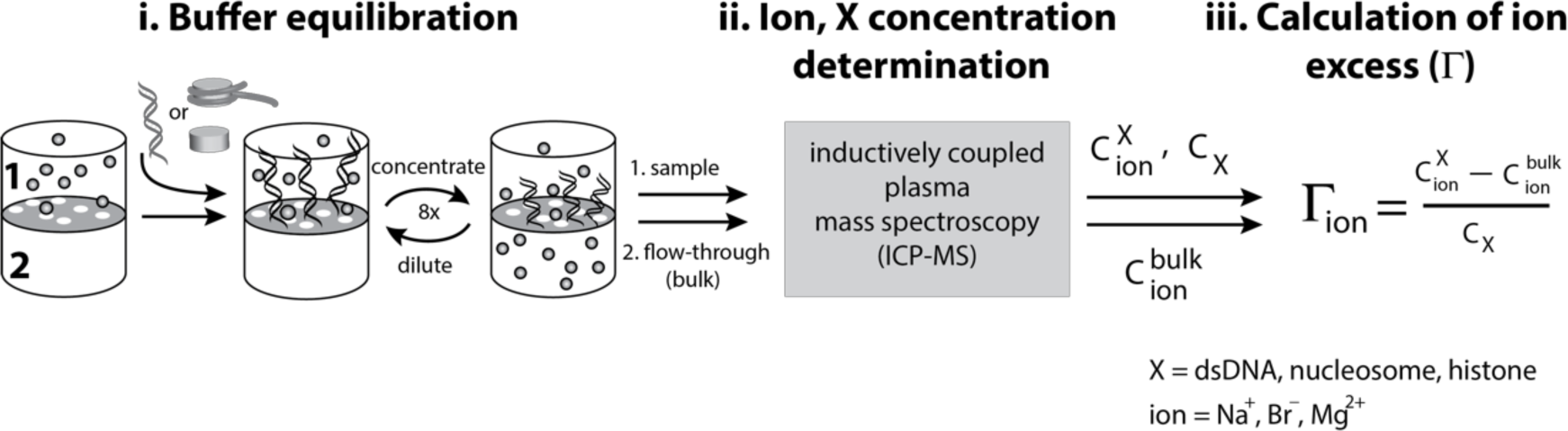
Scheme of ‘ion counting’ to quantify the composition of the ion atmosphere around dsDNA, nucleosomes, histones and the histone octamer.^22, 23, 46^ A description of the ion counting method can be found in the Experimental Method section. Adapted from reference.^23^

In this work, we use ion counting to determine the number of ions associated with free double-stranded (ds)DNA and with nucleosomes, providing a quantitative comparison of their net electrostatic fields. We find that canonical nucleosomes preferentially attract cations (‘counterions’) over anions and do so to an extent similar to non-nucleosomal DNA, confirming Model II prediction of a strong negative potential around nucleosomes. The studies presented herein are foundational for considering the physical and energetic basis for DNA compactions and chromatin organization as well protein binding to nucleosomes and subsequent functional consequences.

## Results

### Strategy to measure nucleosome electrostatics

Polyelectrolytes such as DNA (i.e., a class of macromolecules containing charged groups, either positively, negatively or both) are surrounded by ions that counterbalance their charge. Poisson Boltzmann (PB) electrostatic theory predicts that this charge balance is achieved differently for molecules of low vs. high charge density (Figure 1C). Specifically, weakly charged molecules achieve charge neutrality by equally attracting counterions (i.e., ions with charge opposite to the molecule) and excluding coions (i.e., ions with the same charge to the molecule) (Figure 1C, left). In contrast, molecules with high charge densities and thus with strong electrostatic fields, like DNA, are predicted to achieve charge neutrality by preferentially attracting counterions (cations for DNA) and excluding fewer coions (anions for DNA; Figure 1C, right); the strong electrostatic field of highly charged molecules can counteract the thermal motions of cations resulting in their condensation around the molecules and hence the larger number of cations than anions.^33, 34, 37-39^ This theoretical preference is well established by ion counting experiments.^22, 23, 27, 28^

The degree of preference for counterion attraction varies continuously with charge density, as shown schematically in Figure 1D. Thus, the relative amount of counterion attraction and coion repulsion provides a measure of a molecule’s overall electrostatic field. Specifically, we define β in eq. 1 as the fraction of charge neutralization that arises from association of cation (β_+_) vs. exclusion of anion (β_-_), where *N*_*counterion*_ is the number of attracted counterions, *N*_*coion*_ is the number ofexcluded coions, and *q* _molecule_ is the molecule charge. Because there is overall charge neutrality, the sum of the β values must be one (eq. 1c).

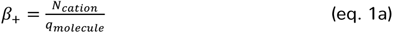

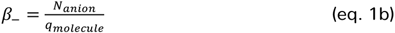

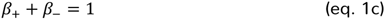

The first method we use to test a molecule’s overall electrostatic potential is direct measurement of β. Model I (Figure 1B), in the extreme, predicts low overall charge density resulting in equal cation attraction and anion repulsion around the nucleosome—i.e., β_+_ = β_-_ ≈ 0.5. Model II predicts β_+_ >> β_-_, with values similar to or more asymmetric than free DNA.

A second way to test a molecule’s overall electrostatic potential is to compare the attraction of counterions of different charge, such as Mg^2+^ vs. Na^+^ for DNA.^24, 25, 40^ PB theory predicts that the preference for divalent over monovalent increases as the strength of the molecule’s electrostatic field increases. For low charge density molecules, the preference for Mg^2+^ over Na^+^ simply follows the bulk composition and ionic strength. For higher charge density molecules, the preference for Mg^2+^ is greater, as each associated Mg^2+^ can interact favorably with multiple closely spaced negative charges when a molecule has high charge density. This preference has also been experimentally verified for DNA.^22, 24, 41-44^ Thus, the Mg^2+^:Na^+^ ratio provides a second measure of the electrostatic character.^25, 44^

### Ion counting reveals a high negative electrostatic potential of nucleosomes

To determine the effect of nucleosome formation on DNA electrostatics we measured the ions associated with free DNA and with nucleosomes by ion counting (Figure 2) and from those values we calculated β_+_ and β? for Na^+^ and Br^—^, respectively. Na^+^ and Br^—^ were chosen for their ease and accuracy of detection by mass spectrometry and because they behave similarly to the more physiological K^+^ and Cl^—^ ions.^23, 24^

Counting ions around 147 bp DNA revealed 5.4-fold preferential attraction of cations with respect to the anion exclusion, giving β_+_ = 0.85 ± 0.02, β_-_ = 0.16 ± 0.01 (Figure 3A – Source Data 1). This large asymmetry in β coefficients is indicative of the strong electrostatic field of dsDNA.

**Figure 3.**
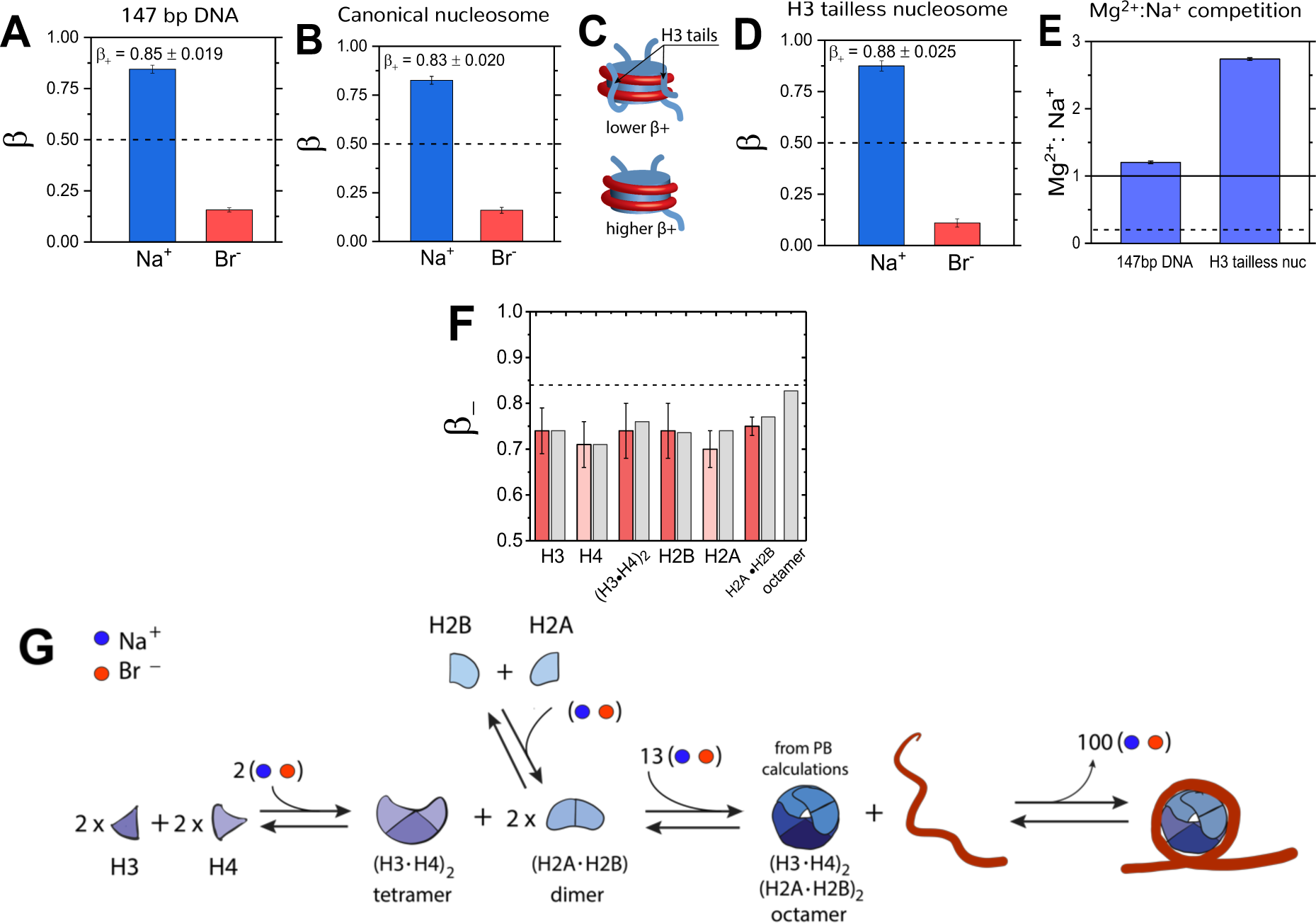
Quantification of electrostatic properties of a free dsDNA, nucleosomes, and histones by ion counting. (A) β coefficients of accumulated Na^+^ cation (blue bar) and excluded Br^-^ anions (red bar) around 147 bp DNA, (B) canonical nucleosomes, (D) H3 tailless nucleosomes. The β_+_ values for the canonical and H3-tailless nucleosomes are significantly different based on two-sample t-test, with p-value = 0.028. Dashed lines represent β = 0.5, the value predicted by Model I. Bulk concentration of NaBr was 10 mM in 2 mM Na-EPPS, pH 7.5 (E) Mg^2+^ vs. Na^+^ competition assay for 147 bp DNA and H3 tailless nucleosome. The solid line represents measurements for a model system, a short 24 bp DNA and the dashed line represents Mg^2+^:Na^+^ ratio for a low charge density molecule, an equivalent to β = 0.5. (F) β_-_ coefficients for histone proteins and their complexes. Experimental results are shown in red bars and theoretical PB calculations are shown in gray bars. Ion counting experiments were carried out at 40 mM NaBr in 2 mM Na-EPPS, pH 7.5, whereas PB calculation were carried out at 40 mM monovalent salt. Dashed line represents the β_+_ value for 147 bp DNA from Fig. 3A. Each data point is the average of 3-8 independent repeats. See Figure 3 – Source Data 1-7 for raw data. (G) Schematic representation of octamer and nucleosome formation. Based on ion counting results and PB calculations for the histone octamer (Figure 3 – Source Data 1-7) we were able to estimate the number of ions required at each step of the assembly, shown by the numbers in the assembly scheme. The formation of the octamer complex requires an uptake of ions to attenuate electrostatic repulsion between positively charged histone proteins, whereas the ion release accompanies nucleosome formation. Note that an equal number of cations and anions are taken up or released, as required (counterintuitively) to maintain charge neutrality (see references ^35, 50, 51^ for explanation).

We carried out the analogous experiment with nucleosomes reconstituted in vitro on the same 147 bp DNA (see Methods). The total charge of the nucleosome (q) from ion counting (see Methods, eq. 3) is considerably lower than that of the 147 bp DNA alone: q_Nuc_ = –144.0 ± 1.7 vs. q_DNA_ = –292.0 ± 4.9 for the nucleosome and the DNA, respectively. This decrease is expected from the association of the DNA to a positively charged histone octamer of total charge +148.0e, as estimated from the amino acid composition (see Figure 3 – Source Data 5) and PDB2PQR calculations.^45^

However, despite the overall reduction in charge by more than two-fold, the excess attraction of cations for nucleosomes remained similar to the dsDNA alone, with β_+_= 0.83 ± 0.020 and β_-_ = 0.17 ± 0.015 based on eight independent determinations (Figure 3B, Figure 3 – figure supplement 1&2, and Figure 3 – Source Data 2). These results provide strong evidence against Model I, which predicts substantial attenuation of the nucleosomal DNA electrostatics and β_+_ close to 0.5.

### Removal of H3 histone tails increases the nucleosome electrostatic potential

The canonical nucleosome is comprised of histone proteins that have N- or C-terminal disordered and mobile extensions, referred to as tails, with a preponderance of positive charge. These tails form regions of positive electrostatic field within nucleosomes and presumably affect the attraction of cations and exclusion of anions. For instance, tails can interact with the nucleosomal DNA and provide electrostatic screening of the DNA charge instead of cations, thereby lowering the electrostatic potential (Figure 3C).^47-49^

To test electrostatic effects of the tail, we reconstituted nucleosomes containing H3 histone proteins lacking their tails (“tailless”; see Methods) and carried out the same analysis as we did for the canonical nucleosomes. The measured overall charge of the H3-tailless nucleosome was more negative (e.g. q=-160 ± 3.1) and in good agreement with theoretical predictions based on the amino acid composition of the H3-tailless histone octamer and the charge of the 147bp DNA (e.g. −160e). Ion counting revealed larger β_+_ and lower β_-_ coefficients compared to values of the canonical nucleosome; β_+_ = 0.88 ± 0.025 and β_-_ = 0.11 ± 0.020, respectively (Figures 3B&D, Figure 3 – Source Data 3). The H3-tailless nucleosome attracts approximately 20% more Na^+^ than the canonical nucleosome. Thus, this result shows that the electrostatic potential around nucleosomes increases when positively charged tails of the H3 histone are removed (Figures 3A–D); the H3-tailless nucleosome on the electrostatic scale (Figure 1E) shifts further to the right. This result confirms that histone tails can make local contacts with the nucleosomal DNA and hence participate in mitigating the negative electrostatic potential of the nucleosomal DNA.

### Mg^2+^ vs. Na^+^ competition substantiates a stronger electrostatic potential of nucleosome compared to free dsDNA

The increased preference for association with divalent over monovalent cations (M^2+^ and M^+^, respectively) as negative charge density increases provides a second measure of macromolecule electrostatics (see ‘Strategy to measure nucleosome electrostatics’ above). M^2+^:M^+^ competition is predicted by PB theory to depend linearly on a molecule’s charge density (ρ) and hence is more sensitive to variations of molecule electrostatics than the β values, which show exponential dependences on ρ and thus have limited ability to resolve electrostatics of molecules with high charge density (see Figure 3E – figure supplement 3).

Given that H3 tails partially counterbalance the negative potential of the nucleosomal DNA, as observed above, we carried out this test with the H3-tailless nucleosomes. We previously measured equal amounts of Mg^2+^ vs. Na^+^ (Mg^2+^: Na^+^ ratio of 0.97 ± 0.06) around a 24 bp DNA despite a bulk concentration ratio of 1 Mg^2+^ per 10 Na^+^ (Figure 3E – Source Data 4).^24^ A weakly charge molecule (Model I) would attract only 1 Mg^2+^ for every 5 Na^+^.

We found that the Mg^2+^: Na^+^ ratio around free 147 bp DNA was 1.15 ± 0.02 (95.0 Mg^2+^ and 81.0 Na^+^; Figure 3 – Source Data 4), very similar to the value obtained for the 24 bp DNA (Figure 3 – Source Data 4), under the same experimental conditions (2.5 mM Mg^2+^ and 25 mM Na^+^). In contrast, for the H3 tailless nucleosomes, the ratio of associated Mg^2+^: Na^+^ was 2.75 ± 0.17, with 58 ± 1.0 Mg^2+^ and 21 ± 1.5 Na^+^ ions (Figure 3E). Thus, the Mg^2+^ vs. Na^+^ competition result provides additional support for Model II—that nucleosomes are highly-charged polyelectrolytes—and also suggests an increase of the electrostatic potentials of the nucleosomal DNA upon complexation into the nucleosome—i.e., that the electrostatic potential of the H3-tailless nucleosome is stronger than the potential of the free 147 bp DNA, although some of the increased Mg^2+^ attraction could arise from direct interactions with nucleosomal DNA, which could reflect DNA rearrangements as well as increases electrostatic potential. The likely origin of the overall high electrostatic potential of nucleosomal DNA is described in the Discussion.

### Histone proteins are positively charged but only partially attenuate the DNA electrostatic potential

How does the overall electrostatic potential of the nucleosome remain highly negative despite of the neutralization of half of the DNA overall charge by the positively charged histone octamer? Two classes of models could explain this: i) the positive charges of histone octamer are broadly dispersed rendering the complex a weak polyelectrolyte and unable to counterbalance the DNA electrostatic field, (i.e., it is like the schematic on left in Figure 1D) or ii) the histone octamer is a strong polyelectrolyte as the DNA (i.e., it is like the schematic on right in Figure 1D) but its positive polyelectrolyte character may be overcome by an increase in the DNA electrostatic field accompanying DNA compaction during nucleosome formation.

To distinguish between these models, we determined the polyelectrolyte character of histone proteins and their stable sub-complexes (e.g. H2A·H2B dimer and (H3·H4)_2_ tetramer), by quantifying their β_+_ and β_–_ values through ion counting. We measured on average 2.6-fold preferential attraction of anions with respect to cation exclusion for histones (e.g. β_+_ = 0.28 ± 0.06 and β_-_ = 0.72 ± 0.04, Figure 3G, Figure 3 – Source Data 6&7) and a small increase of anion attraction and weaker repulsion of cations for the H2A·H2B dimer and (H3·H4)_2_ tetramer (e.g. on average β_+_ = 0.25 ± 0.03 and β_-_ = 0.74 ± 0.01). These results indicate that histone proteins are positively charged and they are not weak polyelectrolytes (i.e., β_-_ > 0.5), yet their polyelectrolyte character is not as strong as dsDNA (β_-_ = 0.72 ± 0.04 vs. β_+_ = 0.85 ± 0.02).

We also determined the electrostatic potential of the histone octamer core. However, as previous work has indicated that the octamer conformation is not stable under physiological or lower salt concentrations in the absence of DNA^52^ and because these conditions are required for ion counting experiments, we could only determine the electrostatic potential of the histone octamer through Poisson-Boltzmann (PB) calculations. To validate this approach, we first compared the experimental and theoretical β_-_ value of anion attraction for individual histone proteins and their stable sub-complexes and observed a good agreement for the predicted and measured values (Figure 3F).

Comparison of the PB calculations suggests that the octamer core attracts more anions than histone proteins alone (predicted β_-_ = 0.84 and β+= 0.17 vs. the predicted average histone β_-_ = 0.73) and the histone octamer is a strong polyelectrolyte, comparable in strength but opposite in field to free dsDNA (i.e., the fraction of the octamer charge neutralization by anions (β_-_ = 0.84) is similar to the fraction of the DNA charge neutralization by cations (β_+_ = 0.85 ± 0.020)). Indeed, our PB calculations suggest that the assembly of histones into octamers increases the electrostatic potential around the octamer and that this process requires an uptake of approximately 13 ions to balance the electrostatic potential increase (Figure 3G). Taken together, our findings raise the important question why histone octamers, despite their strong electrostatic potential (as indicated by PB calculations, Figure 3F), can only partially attenuate the electrostatic charge of DNA in nucleosomes (as evident from our ion counting measurements, Figure 3B-3E). We propose explanations for this phenomenon in the Discussion.

## Discussion

We have carried out the first experimental studies on the ion atmosphere around nucleosomes. Our results provide quantitative insights into the electrostatic properties of nucleosomes and have allowed us to distinguish between two opposing models: Model I, in which nucleosome formation greatly ameliorates the DNA’s electrostatic potential, as presented in standard molecular biology textbooks^11^ (Figure 1B), and Model II, which arises from electrostatic theories and calculations and predicts no reduction of DNA’s electrostatic potential, despite the nearly two-fold decrease in the overall nucleosome charge compared to free DNA (Figure 1B). The observed strong cation association with nucleosomes and preferential association of Mg^2+^ over Na^+^ provide strong experimental support for Model II, raising important questions about both the nature of the strong overall electrostatic potential of nucleosomes and how it affects DNA compaction and chromatin function.

### How does the overall electrostatic potential remain strong, with the net charge of the nucleosome less than that of free DNA?

The simplest explanation for the maintained high electrostatic potential comes from inspecting the nucleosome structure. Approximately, only half of the DNA contacts the histone octamer, and the remaining part is exposed to solution and presumably not subjected to the electrostatic screening from the octamer core (Figure 4A). Thus, one possibility is that the free part of the nucleosomal DNA behaves as a sheath with similar electrostatic properties as the free DNA and define the overall electrostatic character of the nucleosome. However, our ion counting studies on the H3-tailless nucleosome show that the electrostatic potential of the nucleosomal DNA is even stronger compared to the free 147 bp DNA (Figures 3D and 3E), arguing against this simple model.

**Figure 4.**
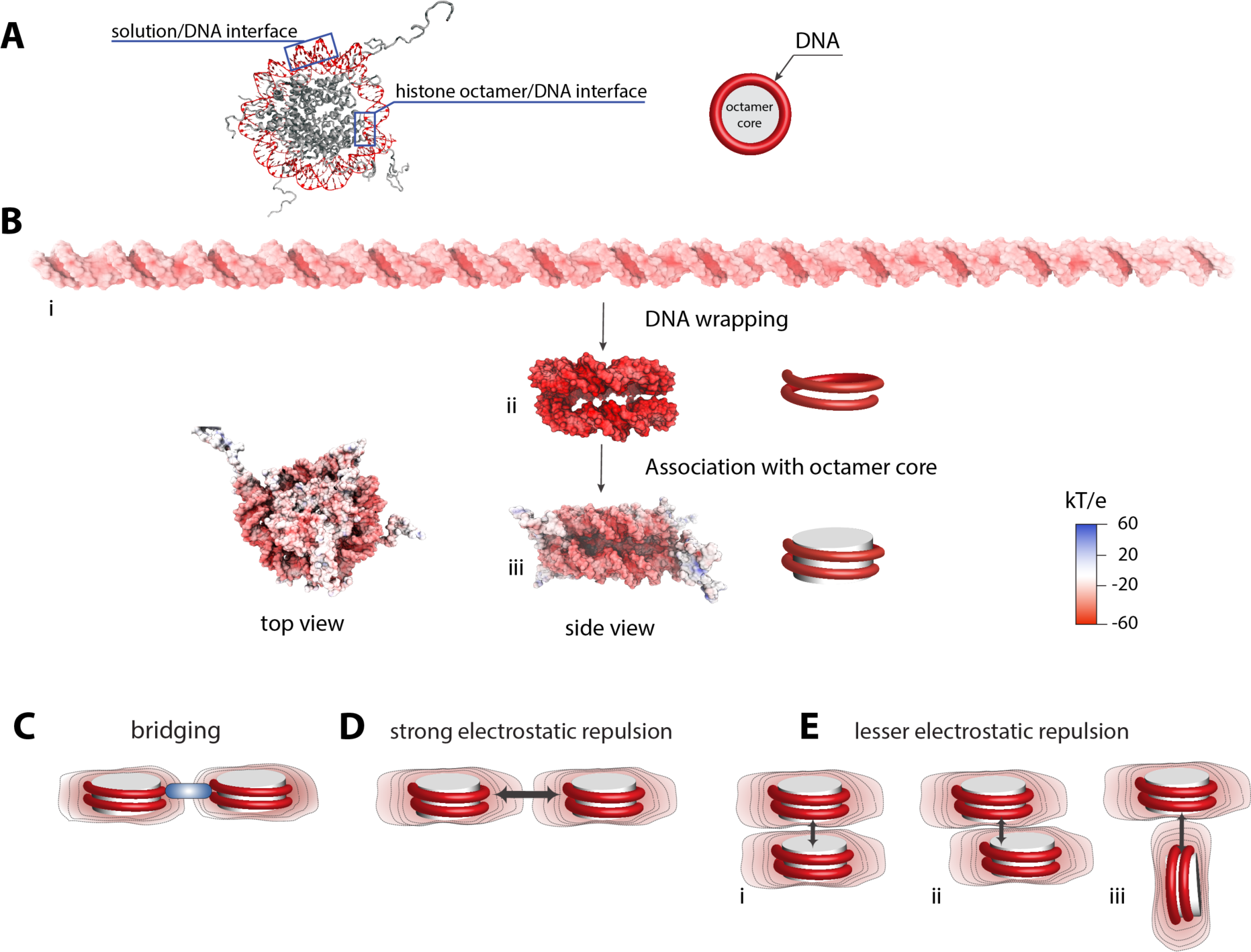
Electrostatic surface potential of a nucleosome and a dsDNA. (A) Crystal structure of a nucleosome (pdb 1kx5).^8^ (B) Poisson-Boltzmann calculations of electrostatic surface potential of a dsDNA and a nucleosome (pdb 1kx5); the electrostatic potential ranges from +60 (blue) to −60 (red) kTe^-1^, which highlights the difference in potential magnitude around the DNA and octamer core. The electrostatic potential mapped to the molecular surfaces was calculated using Adaptive Poisson-Boltzmann Solver (APBS)^68^ and the figures were rendered with VMD.^69^ The electrostatic surface potential of the nucleosomal DNA is largely negative and shown in red. (C)-(E) Schematic representation of electrostatic field lines around a nucleosome and models of internucleosomal interactions. The differences in the electrostatic field as shown in (B) may play an important role in internucleosomal interactions. High electrostatic potential of the nucleosomal DNA disfavors a side-by-side orientation as shown in (D) but introduce the possibility of bridging interactions stabilized by positively charged polyamines and proteins (C, shown in blue). Stacking of nucleosomes with the octamer-to-octamer orientation is energetically more favorable due to weaker electrostatic potentials at the octamer facets as shown in (D).

Electrostatic theory and computational studies provide a more complete model.^13, 14^ The magnitude of the electrostatic potential of a molecule is determined by the molecule’s charge *density* (i.e., the number of charges per given volume, or unit length), rather than the overall charge.^33, 53^ Thus, structural changes that increase the charge density also increase the electrostatic potential, as observed when RNA molecules fold to compact three-dimensional structures.^50, 54, 55^ To illustrate this model, we performed PB calculations (Figure 4B i-iii). For the nucleosome, the wrapping of DNA strands around the histone core brings together backbone phosphoryl groups from distal parts of the DNA helix, and alterations in the duplex geometry decreases the distance between a subset of nearby phosphoryl groups. These features result in a large increase in the negative electrostatic potential (Figure 4B ii). Association of the core only partially mitigates the increased potential, giving a final electrostatic potential that is more negative than free linear dsDNA, in agreement with our experimental data (Figure 4B, i vs. iii). As nucleosomes are stable complexes, there must be a surplus of favorable DNA/nucleosome interactions to overcome, or ‘pay for’, the increased proximity of phosphoryl negative charges and resulting increased electrostatic potential. Our ion counting experiments suggest that part of this energy arises from the significant release of ions accompanying the formation of the nucleosome (Figure 3G), consistent with the difficulty in obtaining equilibrium measurements for nucleosome formation and their extreme sensitivity to solution salt conditions.^9, 56, 57^

### How does the high overall negative electrostatic potential of nucleosomes affect DNA compaction and chromatin function?

Most discussions about the impact of nucleosome formation on DNA focus on the reduced DNA accessibility due to steric occlusion by the histone octamer.^2, 4, 58-61^ However, the electrostatic properties of the nucleosome also strongly influence how DNA interacts with proteins and small molecules and thus how chromatin compacts and functions. Importantly, our ion counting studies offer insights into the underlying mechanisms. Based on the observed stronger preferential association of Mg^2+^ over Na^+^ with the nucleosome compared to the dsDNA, we propose that the higher electrostatic potential and the positioning of phosphoryl oxygen atoms in nucleosomes attracts multivalent counterions (e.g. spermidine and spermine and DNA compacting proteins such as histone H1) substantially more than free dsDNA. These interactions in turn could promote bridging interactions between nucleosomes (Figures 4C) that lead to chromatin compaction. The positively charged histone tails may play similar bridging roles but even more effectively as they are pre-associated with the nucleosome.

Our measurements and calculations also predict the topology type of contacts established in nucleosome arrays. For individual nucleosomes, the strongest negative potential lies on the ‘sides’, so that side-by-side association would be less favored (Figure 4D). In contrast, the negative potential is weakest at the ‘top’, while there is even a weak positive potential in the nucleosome center, where the histone core is located (Figure 4B iii and Figure 4 – figure supplement 1, blue). This difference suggests that there is a strong tendency for nucleosomes to ‘stack’ off-center and to associate perpendicularly, which would align regions of negative and positive potential (Figure 4E). Intriguingly, recent cryo-electron micrographs revealed preferential nucleosome association consistent with these electrostatic precepts.^62^

Apart from such ordered interactions between nucleosomes, dynamic multivalent interactions have recently been implicated in heterochromatin formation by compacting nucleosome arrays into phase-separated, higher-order condensates.^63, 64^ Given the ability of nucleosomes to make strong electrostatic interactions with multivalent counterions demonstrated by our experiments and the long-range nature of these interactions, we hypothesize that nucleosome electrostatics also play a fundamental role in chromatin phase separation.

Furthermore, the non-uniform and concentrated electrostatic potential around the nucleosomal DNA likely not only plays an important role in organizing chromatin, but also in coordinating nucleosome-protein interactions that are at the heart of biological processes like gene transcription or DNA repair. Most DNA binding proteins are positively charged and their relative affinities are expected to be dependent on the local electrostatics of the nucleosome.^15-17, 65-67^ The strong and varied electrostatics of nucleosomes thus introduce an additional variable that Nature has likely utilized in controlling gene expression. Dissecting this remains an important goal for future studies to fully and deeply understand the regulation and misregulation of gene expression.

## Material and Methods

### Reagents

DNA molecules to assembly 147 bp DNA^70^ were purchased from IDT (Ultramer^®^ DNA Oligonucleotide, Integrated DNA Technologies, USA). The purify of DNA (>96%) was verified by 5% native-PAGE gel with the load of 100-200 ng of DNA was per lane, stained with SYBR_™_Gold (Invitrogen, USA) with a DNA detection limit of 25 pg; CLIQS (Totallabs, UK) imaging analysis software was used for gel analysis. Histone expression plasmids were from Narlikar lab and BL21(DE3)pLysS competent *E.coli* cells were from Agilent Technology (USA). All salts were of the highest purity (TraceSELECT^®^ or BioXtra, Sigma-Aldrich USA). All solutions were prepared in high purity water, ultra-low TOC biological grade (Aqua Solutions, USA).

### Protein expression, purification and octamer assembly

All histones from *Xenopus leavis* (H2A, H2B, H4, H3 and tailless H3) were expressed from *E. coli* and purified following published protocols.^71-73^ The tailless H3 histone lacks the N-terminal region of the canonical H3 histone (25 residues including 8 positively charged residues). Purification was carried out by an anion exchange through 5-ml HiTrap Q column followed by a cation exchange through 5-ml HiTrap S HP column. Subsequently, histones were subjected to gel filtration on a Superdex 75 column, to attain high purity. All columns were from GE Healthcare Life Sciences (USA). Histone octamer was assembled from purified histones as described.^71-73^

### Nucleosome assembly

The 147 bp DNA was assembled from equimolar complementary strands (0.1-0.5 mM) in 100 mM Na-EPPS (sodium 4-(2-hydroxyehyl)piperazine-1-propanesulfonic acid), pH 7.5. Samples were incubated at 90 °C for 2 min and gradually cooled down to ambient temperature over 1 h. Non-denaturing polyacrylamide gel electrophoresis showed no detectable single stranded DNA in samples, corresponding to >90% duplex; DNA stained by SybrGold (Invitrogen). Nucleosomes were assembled using published gradient dialysis-based protocols^71-73^ and the purification of the nucleosomes was carried out on a 10 to 30% glycerol gradient. Subsequently, a fraction of collected nucleosomes was loaded onto 5% native-PAGE gel: the amount of nucleosome complex corresponded to > 95% (DNA stained by SYBR_™_Gold)

### Buffer Equilibration-Inductively Coupled Plasma Mass Spectroscopy (BE-ICP MS)

Buffer equilibration for nucleosomes and proteins was carried out following previous procedures.^22, 23^ NaBr and MgBr_2_ samples were prepared in 2 mM Na-EPPS, pH 7.5 and their concentrations were determined by ICP MS. 500 uL–samples of nucleosome (4-12 *μ* M) or proteins (4-100 *μ* M) with the salt of interest were spun down to *Γ* 100 µL at 7000 x g in Amicon Ultracel-30K filters (Millipore, MA) at 4°C (Figure 2). Buffer equilibration was carried out until the ion concentration in the flow-through samples matched the ion concentration in the buffered solution used for the buffer exchanged.^22, 23^ No loss of the nucleosomes or proteins was observed during this procedure; no DNA or proteins were detected in flow-through samples, as determined by ICP MS, assaying the phosphorus content, or UV measuring absorbance at 280 nm. Nucleosomes were intact after the course of ion counting experiments as indicated by non-denaturation PAGE (Figure 3 – figure supplement 2).

### Ion counting

Inductively coupled plasma mass spectrometry (ICP-MS) measurements were carried out using a XSERIES 2 ICP-MS (Thermo Scientific, USA). Herein, ion counting measurements were carried out with bromide salts, as the detection of Br^−^ anion by ICP MS has highest accuracy and precision compared to other halogens. Aliquots (10–20 µL) of nucleosome- or histone-containing sample, the flow-through from the final equilibration, and the equilibration buffer were diluted to 5 mL in 15 mL Falcon tubes with water. Dilution factors, the ratio of diluted to total sample volume, were used to maintain sample concentrations within the linear dynamic range of detection.^22, 23^ Calibrations were carried out using standards from SpexCertiPrep (USA). Quality control samples, containing each element of interest at 100 µM, were assayed every ten samples to estimate measurement precision. To minimize memory effects in Br^−^ detection, a solution of 5% ammonium hydroxide in highly pure, ion-free water (Mili Q) was used as a wash-out solution between measurements.^74^

The count of associated ions around 147 bp DNA and nucleosomes is reported here as a preferential ion interaction coefficient *Γ*_*i*_ (e.g. the number of associated ions, i =+ or -, indicating cation or anion, respectively).^75^ The *Γ*_*i*_ was calculated as the difference in the ion concentration between the equilibrated samples containingdsDNA 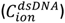, nucleosome 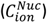, or histones 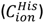 and the bulk solution 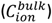, divided by the concentration of the molecules determined by phosphorous measurements using ICP MS (for the DNA and nucleosome) or determined by absorption at 280 nm for histones (eq 2).

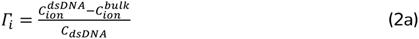

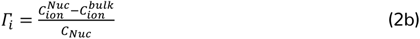

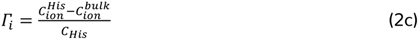

For negatively charged molecules (e.g. DNA and nucleosomes), the cation preferential interaction coefficient, *Γ*_*+*_, is expected to be greater than zero, indicating their accumulation around the negatively charged polyelectrolytes, and *Γ*_*-*_ for an anion is expected to be less than zero due to repulsive interactions with the DNA.

The total charge of the ionic species around molecules was calculated as the sum of the number of ions multiplied by their charge (*z*_*i*_) and must counterbalance the molecule charge (q) (eq. 3).

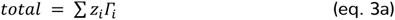

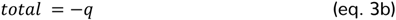

For each ion counting data point reported, at least three measurements were made on three different days with independently prepared samples. Errors are the standard deviation of all measurements

### Poisson Boltzmann (PB) calculations

NLPB calculations were carried out for a 147 bp DNA duplex and nucleosome. The B-form 147 bp DNA duplex was constructed with the Nucleic Acid Builder (NAB) package version 1.5.^76^ PB calculation on nucleosome were carried out using X-ray crystal structure (pdb: 1kx5).^8^ Charges were assigned using the PDB2PQR routine with the Amber parameter.^68^

NLPB calculations were carried out using the Adaptive Poisson-Boltzmann Solver (APBS)^68^ on a 808 × 808 × 862 Å^3^ grid with a grid spacing of 1.8 Å. The ion size equal 2 Å, the simulation temperature was set to 298.15 K and the dielectric constant of the solvent was set to 78. The internal dielectric was set to 2. The solvent-excluded volume of a molecule was defined with a solvent probe radius of 1.4 Å. Boundary conditions were obtained by Debye-Hückel approximation.

The preferential interaction coefficient of ions of valence associated with macromolecules was computed by integrating the excess ion density:^22, 41, 77, 78^

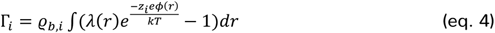

where б_*b*, *ii*_ is the bulk ion density, *λ*(r) is an accessibility factor that defines the region in space that are accessible to ions where *λ*(r)=1 and for the solvent-excluded region –i.e., inside the macromolecule where *λ*(r)=0, *z*_*i*_ is the elementary charge, *ø* (*ri* is the electrostatic potential, *k* is the Boltzmann constant, and *T* is the temperature. The integration volume was defined as the entire volume of a simulation box including the solvent-excluded region in the molecule interior. This approach matches the conditions for the experimental measurement.^23^ Numerical integration of eq. 4 was carried out using a custom written routine in C++, which is available from the authors upon request.

## Acknowledgment

The authors thank Bradley French and Broder Schmidt, members of the Herschlag and Straight lab for helpful discussions and critical advice. The authors thank Guangchao Li from Environmental Measurements Facility at Stanford University for outstanding technical assistance with ICP MS measurements.

## Additional information

### Author ORCIDs

Decision letter and Author response

### Author contributions

Conceptualization: MG and DH; performing experiments: MG; formal analysis: MG; histone plasmids were provided by GN; SJ trained MG how to reconstitute mononucleosomes; writing-original draft: MG, DH; writing—final draft: MG, JS GN, DH; funding acquisition: DH

## Additional files

Supplementary files

- Transparent reporting form:

Data availability

